# Nanopore Sequencing Unveils Somatic Structural Variations as Biomarkers in Laryngeal squamous cell carcinoma Genomes

**DOI:** 10.1101/2025.06.12.659252

**Authors:** Xuyan Liu, Lin Xia, Yixin Qiao, Yang Li, Yan Huang, Bingyan Yue, Xi Liang, Xin Yang, Honghui Zhang, Jiaxun Zhang, Xiao Chen, Dan Xie, Jifeng Liu

## Abstract

Laryngeal squamous cell carcinoma (LSCC) is an aggressive cancer with poor quality of life, largely due to the absence of reliable molecular biomarkers for early diagnosis and prognosis. While somatic structural variations (SVs) have been documented in LSCC, conventional short-read sequencing approaches are inherently limited in detecting of large-scale somatic SVs, which are critical to understanding tumorigenesis. Here, we presented SomaGauss-SV, a somatic SVs detection workflow leveraging nanopore long-read sequencing data. Benchmarking against five paired tumor cell line datasets showed SomaGauss-SV consistently achieves a balanced high precision and recall. SomaGauss-SV applied to 15 paired LSCC tumor-blood samples uncovered a comprehensive SVs landscape and a significant positive correlation between somatic deletion burden and smoking intensity. Furthermore, a high-frequency somatic simple repeat expansion was identified in 20/27 (74.1%) of LSCC patients, upregulating the expression of genes *TP53BP2* and *FBXO28* through spatial proximity. These findings underscore the potential of long-read sequencing and SomaGauss-SV in uncovering previously unexplored genomic rearrangements in LSCC, providing valuable resources for biomarker discovery.

## Introduction

Laryngeal squamous cell carcinoma (LSCC) is the second most common subtype of head and neck squamous cell carcinoma (HNSCC), characterized by marked gender differences and regional variability in both incidence and mortality rates. The incidence rate in males is approximately eight times that of females(Megwalu and Sikora 2014). LSCC pathogenesis is closely related to modifiable risk factors, such as tobacco exposure and alcohol consumption(Bray et al. 2024; Sung et al. 2021; Chen et al. 2024; Johnson et al. 2020). However, the molecular mechanisms by which these exogenous factors induce genomic instability remain unclear. Early symptoms of LSCC are often overlooked, resulting in more than 50% of patients being diagnosed at advanced stages, particularly those with supraglottic tumors(Steuer et al. 2017; Nocini et al. 2020). This delay in diagnosis typically requires aggressive treatments that severely impair patients’ respiratory and vocal functions, thereby significantly dimishing their quality of life(R et al. 2019). Despite advances in treatment regimens, combining surgery with radiotherapy and chemotherapy, the five-year overall survival rate has shown limit improvement(Cavaliere et al. 2021; Wang et al. 2020), highlighting the urgent need for molecular biomarkers specific to LSCC. Compared to other HNSCC subtypes, LSCC has the unique biological characteristics and epidemiological features(Cavaliere et al. 2021; Pan et al. 2025), making specialized research essential to address its unique diagnostic challenges and the critical need for organ function preservation.

In recent years, pan-cancer genomic studies have highlighted the pivotal role of structural variants in the occurrence and progression of cancer(Cosenza et al. 2022). Structural variants (SVs), which encompass large genomic rearrangements exceeding 50 bp, include deletions (DEL), insertions (INS), duplications (DUP), inversions (INV), and translocations (TRA))(van Belzen et al. 2021). These variants drive oncogenesis through mechanisms such as gene disruption and fusion, copy number alterations, and three-dimensional genome reorganization(Beroukhim et al. 2016). Large-scale cancer cohort studies have shown that somatic SVs are the most prevalent class of driver mutations in cancer, vastly outnumbering single nucleotide variants (SNVs) and small insertions/deletions (indels)(Cosenza et al. 2022). Given their abundance and diverse roles in cancer genomes, a comprehensive understanding of somatic SV’s scope and mechanisms is eseential for uncovering key cancer mutations in patient tumors and identifying biomarkers for diagnosis and treatment.

However, despite the widespread clinical application of somatic SV detection methods based on second-generation short-read sequencing, these approaches have inherent limitations in resolving large-scale structural variants, repetitive sequence variants, and complex genomic SVs(ICGC/TCGA Pan-Cancer Analysis of Whole Genomes Consortium 2020). The advent of nanopore long-read sequencing technology offers a promising solution to overcome this bottleneck. With read length exceeding 10 kb, nanopore sequencing can span complex SVs breakpoint regions in their entirety, enabling comprehensive mapping of somatic SV landscapes. Recently, several somatic SVs algorithms, such as nanomonsv(Shiraishi et al. 2023), Severus(Keskus et al. 2025), and SVision-pro(Sahlin et al. 2023), have been developed to harness this potential. However, existing tools still face challenge in correcting alignment biases in repetitive regions, distinguishing mixed signals of multi-scale insertions (INSs) and deletions (DELs), and their practical application in clinical samples of sufficient scale remains limited(Sahlin et al. 2023). Therefore, developing detection workflows suitable for different research scenarios is a key step for advancing the clinical interpretation of SVs.

To this end, this study has developed a somatic SVs detection workflow, SomaGauss-SV, based on nanopore long-read sequencing data, and evaluated its performance using five pairs of tumor-normal cell lines. The results demonstrated that SomaGauss-SV outperforms all other compared somatic SV detection tools, achieving the highest average F1 score, indicating superior balance between sensitivity and accuracy—unlike other tools that often compromise one for the other. Moreover, SomaGauss-SV showed a high overlap rate of 92.2% with the results from other somatic detection software, further confirming the reliability of its findings. Subsequently, using nanopore long-read whole-genome sequencing technology combined with the SomaGauss-SV workflow, we identified somatic SV maps in primary tumors and paired samples from 15 patients with LSCC and performed feature analysis on the variant sequences and regions. We found a significant positive correlation between tobacco exposure and somatic DEL burden. In addition, by incorporating paired samples from an additional 12 LSCC patients, we identified a simple repeat expansion (SRE) that occured in 74.1% (20/27) of LSCC cases. Multi-omics data analysis suggested that this SRE may regulates the expression of genes *TP53BP2* and *FBXO28* through spatial proximity. This hypothesis was further validated through cell experiments. Overall, the SomaGauss-SV developed in this study demonstrates significant advantages in somatic SVs detection, expanding insights into the molecular mechanisms of LSCC and the discovery of potential clinical diagnostic biomarkers. It also provides a wealth of highly promising somatic SVs candidates for future research.

## Results

### SomaGauss-SV demonstrates excellent performance in detecting somatic structural variations

Current long-read sequencing algorithms for detecting SVs often exhibit reduced accuracy in repetitive genomic regions. This limitation arises from inherent biases in traditional seed-chain alignment methods(Sahlin et al. 2023) (Supplemental Fig. S1). The lack of systematic alignment error correction in existing SVs detection tools further constrains their utility for somatic SVs calling. To address these challenges, we present SomaGauss-SV, a novel workflow (Fig. 1A) that integrates a mixed Gaussian distribution model with alignment error correction, specifically designed to improve the accuracy of somatic SVs detection.

**Fig. 1.**
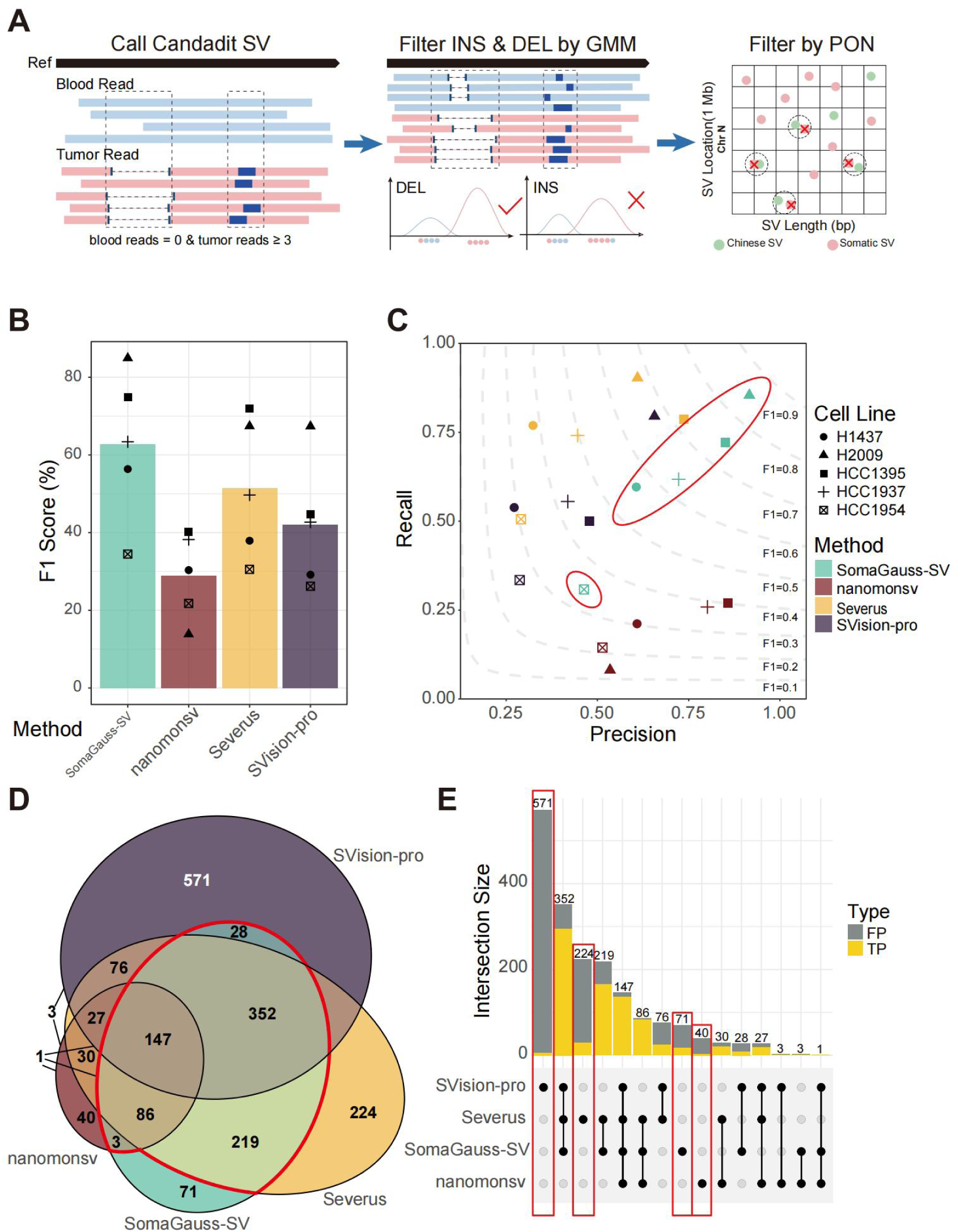
**Overview of somatic SV detection softwares evaluation. (A).**Schematic of the SomaGauss-SV workflow, which consists of three main steps: (1) detection of candidate somatic SVs, screening of somatic SVs through a merge method, (2)filtering of insertions (INS) and deletions (DEL) using a Gaussian Mixture Model (GMM), and (3) filtering of SVs using data from the Chinese normal population. **(B).**Bar chart shwoing the F1 scores for different software. The height of the bars represents the average F1 score for each software, and different shapes of points indicate the F1 scores for different cell lines. **(C).**Contour plot of software evaluation. The X-axis represents precision, the Y-axis represents recall, the contours in the plot represent F1 scores, and software and cell lines are distinguished by the color and shape of the points, respectively. **(D).**Venn diagram of software detection results in the HCC1395 cell line. Numbers indicate the number of SVs in the intersections between software, and red lines indicate the intersections involving SomaGauss-SV with other software. **(E).**UPSet plot of true positive (TP) proportions for different software in the HCC1395 cell line. Yellow indicates TP, gray indicates false positives (FP), red boxes indicate the proportion of TP unique to each software, and black circles below indicate intersections between software.

The SomaGauss-SV workflow consists of three primary analytical phases (Fig. 1B). First, we conduct preliminary somatic SVs detection using Sniffles2 on merged tumor/normal alignment data, retaining candidate somatic SVs supported exclusively by tumor-derived reads (at least three supporting reads). In the second phase, we apply differential filtering strategies: (i) for candidate somatic deletions (DELs), we use Gaussian mixture modeling in conjunction with statistical testing to characterize deletion length distributions; (ii) for putative insertions (INS) and duplications (DUP), we utilized Straglr(Chiu et al. 2021), a length-based Gaussian modeling approach with proven efficacy in repetitive regions. Finally, to minimize the risk of germline SVs contamination from sequencing depth artifacts, we employ Jasmine(Kirsche et al. 2023) for systematic filtering against a panel of normals (PON) derived from the Chinese normal population SVs datasets(Wu et al. 2021), yielding high-confidence somatic SVs calls (Methods).

To systematically assess the performance of SomaGauss-SV in somatic SVs detection, we utilized a publicly available benchmarking resource comprising five tumor-normal cell line pairs: HCC1395/HCC1395BL, HCC1937/HCC1937BL, HCC1954/HCC1954BL, H1437/H1437BL, and H2009/H2009BL(Keskus et al. 2025). These paired samples were sequenced using Oxford Nanopore technology R10 sequencing, achieving a blood median coverage of 28X (range: 24X-47X) and a tumor median coverage of 79X (range: 75X-110X), with read median N50 values of 37kb for blood and 28kb for tumor (Supplementary Table 1). In this comparative analysis, we evaluated SomaGauss-SV alongside established somatic SVs callers, including nanomonsv(Shiraishi et al. 2023), SVision-pro(Wang et al. 2025), and Severus(Keskus et al. 2025), with performance benchmarked against the curated SVs ensemble from this resource. To ensure methodological consistency across the comparison groups, we deliberately omitted the PON filtration step, given the controlled nature of the cell line analyses.

SomaGauss-SV consistently produced the highest F1 scores across all 5 cell line pairs, achieving a mean F1-score of 62.8% (±19.2% SD), significantly outperforming Severus (51.5% ±18.0%), SVision-pro (42.0% ±16.3%), and nanomonsv (28.9% ±11.1%) (Fig. 1B). While all tools exhibited variability across cell lines with distinct SVs class compositions (Supplemental Fig. S2A). Notably, SomaGauss-SV achieved peak performance (84.9% F1-score) in the H2009 cell line, which is characterized by insertion-dominant SVs profiles (mean comparator performance: 57.3%, Supplemental Fig. S2A), highlighting its particular strength in resolving this challenging variant class. In contrast, the HCC1395 cell line, with its high SVs burden, showed narrower performance differentials between SomaGauss-SV (74.9%) and Severus (72.0%), both substantially outperforming the other tools (mean: 42.5%). Interestingly, the genome-doubled HCC1954 cell line exhibited universally lower F1-scores across all platforms (mean: 28.2% vs. 50.8% in non-polyploid lines), suggesting that polyploidization introduces systematic challenges in SVs detection. A critical analysis revealed the precision-recall trade-offs of each tool (Fig. 1C): nanomonsv exhibited higher precision (59.1% vs. competitors’ mean precision 50.7%), while Severus and SVision-pro prioritized recall (64.3% vs. competitors’ mean recall 40.6%). SomaGauss-SV uniquely balanced both metrics (mean precision: 64.0%, mean recall: 61.9%). This indicated that SomaGauss-SV is a well-rounded tool for somatic SVs detection, while other software may be more suitable for specific scenarios depending on whether precision or recall is more important.

The inter-cell line performance variation strongly correlated with the composition of somatic SVs types. To explore this relationship further, we conducted stratified analyses across SVs types using type-specific ensembles (Supplemental Fig. S2B). SomaGauss-SV consistently outperformed other tools across all SVs types. Notably, in INS detection, SomaGauss-SV achieved an F1-score of 59.3%, outperforming comparator tools by 13.9-93.1 % (mean F1-score of comparators: 25.5%). For DUP, SomaGauss-SV exhibited performance comparable to Severus (67.7% vs. 69.1%). While all tools showed relatively higher efficacy in DEL detection (mean F1-scores: 45.4-70.2%), both breakend (BND) and inversion (INV) detection proved challenging across platforms (mean F1 < 5.7%). This mechanistic understanding explains the universally depressed performance in the HCC1954 cell line, where BNDs constituted 64.2% of SVs (Supplemental Fig. S2B), highlighting critical gaps in current SV detection paradigms for complex rearrangements. Given these limitations, our clinical pipeline incorporating SomaGauss-SV includes a mandatory manual curation step for INV and BND calls in tumor-blood paired samples, ensuring high accuracy in detecting these two somatic SVs types (Methods).

Current benchmarking paradigms for somatic SVs detection rely on consensus approaches integrating multiple sequencing platforms and analytical tools(Shiraishi et al. 2023; Wang et al. 2025; Keskus et al. 2025). Using the the widely adopted HCC1395/HCC1395BL cell line pair(Shiraishi et al. 2023; Wang et al. 2025; Keskus et al. 2025), we systematically evaluated the reliability of SVs callsets through inter-tool concordance analysis (Fig. 1D). SomaGauss-SV demonstrated superior consensus validity, with 92.2% overlap across comparator tools, outperforming nanomonsv (88.1%), while maintaining the highest true positive rate (23.94%) among its 71 unique calls (Fig. 1E). Notably, although the intersection of somatic SVs callers improved true positive proportions (single-tool maximum SomaGauss-SV: 78.6% vs 4 tools consensus: 92.5%), this strategy resulted in substantial variant loss (52.3%) and introduced systematic class biases (e.g., the proportion of DEL detection increased by 29.0%; Supplemental Fig. S2C), underscoring the limitations of consensus-based filtering for comprehensive SV profiling. Collectively, these findings position SomaGauss-SV as the optimal solution for our LSCC study, balancing precision and recall while preserving both high-confidence consensus variants and biologically relevant unique calls - critical advantages for downstream clinical interpretation.

### Pathological information and sequencing quality data display of ONT WGS samples

To interrogate the somatic SVs landscape of LSCC, we included tumor samples, comprising with mateched blood samples from 15 treatment-naive primary LSCC patients (Fig. 2A). These tumor samples represented a broad spectrum of clinical stages and anatomical subtypes (Fig. 2B and 2C). All patients were male, with a mean age of 63 years (median: 60). Smoking exposure was assessed using the smoking index, calculated as the number of cigarettes per day multiplied by years of smoking(Zhou et al. 2023). 14/15 patients had a history of smoking, with an average smoking index of 648 (median: 610) (Supplementary Table 2, Fig. 2D). These cohort characteristics align with previous studies, demonstrating that LSCC predominantly affects middle-aged and elderly males and is strongly associated with smoking(Johnson et al. 2020; Lechien et al. 2022), reflecting the typical demographic profile of LSCC patients.

**Fig. 2.**
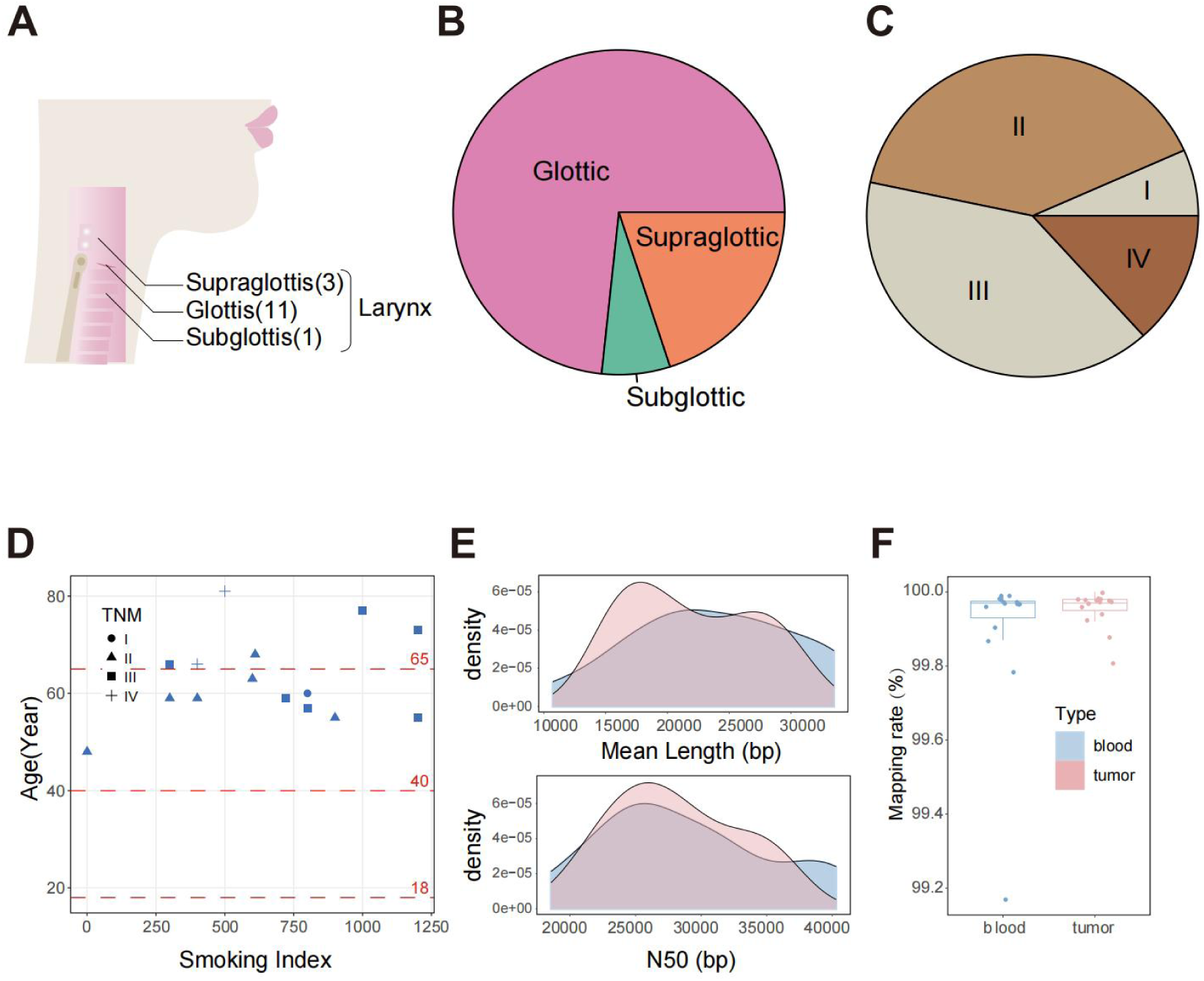
**Overview of pathological information and sequencing data. (A).**Display of sampling sites for LSCC. There are 3 cases of supraglottic type, 11 cases of glottic type, and 1 case of subglottic type. **(B).**Pie chart of the proportion of sampling sites for LSCC. **(C).**Pie chart of the proportion of TNM staging for LSCC patients. **(D).**Pathological scatter plot for LSCC patients. The X-axis represents the smoking index of patients (calculated as: number of cigarettes per day × smoking years), the Y-axis represents patient age, and the shape of the points indicates TNM staging. **(E).**Reads length distribution plot of nanopore sequencing. The upper part shows the distribution of average read lengths (bp) for tumor (pink) and blood (blue), and the lower part shows the N50 distribution for tumor and blood. **(F).**Box plot of read Mapping rates. Blue represents blood, and pink represents tumor.

Whole-genome sequencing of paired tumor and blood samples was performed using the PromethION platform (R9: 9 samples, R10: 6 samples; Supplementary Table 2), generating 2.10 Tb of data. After quality control (sequencing quality > 7, read length > 1 kb), tumor samples yielded an average N50 of 28 kb (range: 20 kb to 36 kb) and an average of 94 billion bases per sample, equating to a mean sequencing depth of 30X (range: 28X to 33X, Fig. 2E). Matched blood samples generated an average N50 of 28 kb (range: 19 kb to 40 kb) and an average of 49 billion bases per sample, resulting in a mean sequencing depth of 15X (range: 12X to 18X, Fig. 2E). Alignment of the clean long-read sequencing data to the hg38 reference genome using minimap2 yeilded average alignment rate of 99.95% for tumor samples and 99.90% for blood samples (Supplementary Table 3, Fig. 2F).

### Discovery of the positive relation between soamtic DEL burden and tobacco exposure

Using SomaGauss-SV, 8128 somatic SVs were identified across the 15 paired samples, with an average of 541 somatic SVs per tumor samples. Somatic insertions (INS) were the most common variant, comprising 54.7% of the total, followed by deletions (DEL, 19.7%), rearrangements (BND, 18.0%), inversions (INV, 4.0%), and duplications (DUP, 3.6%) (Supplemental Fig. S3A). The distribution of SV types in individual samples was consistent with this overall pattern (Fig. 3A). Notably, samples with higher smoking indices displayed a significantly greater somatic SV burden (Fig. 3A) (Pearson correlation coefficient, p<0.05, r=0.54; Supplemental Fig. S3B). In contrast, no significant correlations were observed between smoking index and age (Pearson correlation coefficient, p>0.05, r=0.29, Supplemental Fig. S3C) or TNM stage (Student’s t-test, p>0.05) (Supplemental Fig. S3D). Further analysis revealed a significant positive correlation between smoking index and somatic DEL burden (Pearson correlation coefficient, p<0.05, r=0.67, Fig. 3B), while no significant correlations for other SV types (Supplemental Fig. S3E). This finding similar with previous study indicating that smoking increases the somatic mutation burden in bronchial epithelial cells and LSCC(Yoshida et al. 2020; Degawa et al. 1994), suggesting that tobacco exposure may promote somatic DELs by introducing DNA double-strand breaks(Yoshida et al. 2020; Degawa et al. 1994; Aw et al. 2021).

**Fig. 3.**
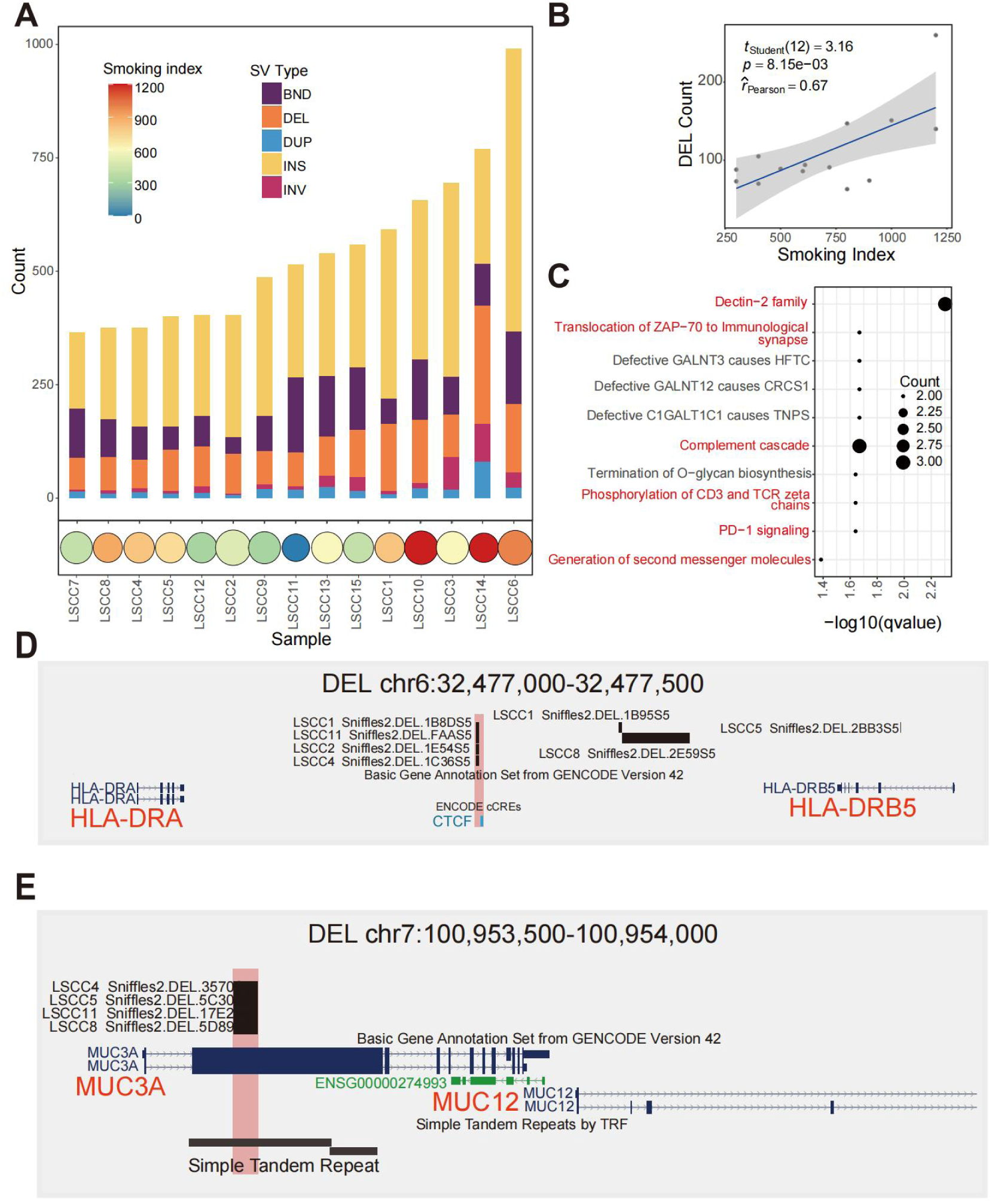
**Relationship between somatic DELs and tobacco exposure. (A).**Stacked bar chart of somatic structural variation counts and smoking index distribution. The upper stacked bar chart shows the number of different SVs for each sample, and the lower bubble chart shows the smoking index for corresponding samples, where higher indices are represented by redder colors. (**B).**Correlation plot of somatic DEL counts and smoking index. The X-axis represents the smoking index, and the Y-axis represents the number of DELs, with degrees of freedom, P-value, and Pearson correlation coefficient marked on the plot. **(C).**Reactome pathway enrichment plot for genes affected by common DELs in multiple samples from smokers. The X-axis represents -log10(q-value), and the Y-axis represents the names of enriched pathways, with immune-related pathways highlighted in red. The size of the circles indicates the number of enriched genes. **(D).**Browser plot of somatic DEL (chr6:32,477,000-32,477,500) illustrating the following epigenomic signals: sample DEL location, ENCODE-annotated gene transcript, and Candidate Cis-Regulatory Elements (cCREs) predicted by ENCODE. **(E).**Browser plot of somatic DEL (chr7:100,953,500-100,954,000) illustrating the following epigenomic signals: sample DEL location, ENCODE-annotated gene transcript, and Simple Tandem Repeats identified by TRF.

To investigate the functional impact of somatic DELs in smoking LSCC, we next focused on the 220 somatic DEL regions shared by more than three smoking-related LSCC samples. GREAT(McLean et al. 2010) mapped 211/220 loci to 49 putative distal regulatory target genes. Pathway enrichment analysis revealed that six of the top ten enriched pathways were immune-related, including Dectin-2 family(Graham and Brown 2009), Translocation of ZAP-70 to Immunological Synapse(Blanchard et al. 2002), and Complement Cascade(Afshar-Kharghan 2017)(Fig. 3C). GO analysis further supported this immune-focused pattern, with nine of the top ten biological processes related to lymphocyte activation and cytokine responses (Supplemental Fig. S3F). These findings supported the role of smoking in tumorigenesis through immune-inflammatory responses(Aw et al. 2021), suggesting that somatic DELs frequently carried by high-smoking LSCC patients may regulate the smoking-associated immune microenvironment.

Among the immune-related somatic DEL regions, a particular a somatic deletion (DEL), shared by four smoking-related LSCC tumors, may affect the expression of *HLA-DRA* and *HLA-DRB5*, which are key immune modulators involved in tumor immune evasion. This high-frequency DEL is located near a CTCF binding site, identified in the MCF-7 and A673 cell lines, 265 bp apart (Fig. 3D). The CTCF site is predicted to regulate *HLA-DRA* and *HLA-DRB5*(ENCODE Project Consortium 2012). We propose that the somatic DELs disrupt CTCF binding, leading to altered chromatin architecture and dysregulated expression of *HLA-DRA* and *HLA-DRB5*. Additionally, we identified a somatic DEL in the *MUC3A* gene, which encodes mucin and has been associated with smoking-induced changes in chronic obstructive pulmonary disease (COPD)(Merikallio et al. 2023). *MUC3A* expression is reported to be higher in smokers than in non-smokers due to tobacco exposure(Gendler and Spicer 1995). The somatic DEL, located at exon 2 of *MUC3A* gene (1120 bp to 1123 bp), contains a simple tandem repeat sequence (motif: ACC; Fig. 3E). This exon encodes a peptide rich in threonine (Thr) and serine (Ser), which are essential for glycosylation and for mucin 3A’s role in protecting the epithelial barrier from environmental damage(Bhatia et al. 2019; Kufe 2009). The somatic DEL might lead to truncation of this peptide, which may impair mucin 3A function .

### Charateristics of Somatic SVs

We characterized somatic SVs in LSCC by analyzing SV size, sequence context, and breakpoint distribution. Our results highlight distinct length preferences across SV types (Fig. 4A). Somatic DELs and INSs were characterized by shorter lengths and narrower distributions. DELs (mean: 1493 bp, range: 50 bp to 30 kb), had a peak at size of 70 bp, while INSs (mean 1107 bp, range: 50 bp to 24 kb), exhibiting a tri-modal distribution with peaks at 70 bp, 170 bp, and 300 bp. In contrast, DUPs and INVs displayed larger, more dispersed length distributions. DUPs (mean: 39 kb, range: 121 bp to 3.9 Mb), showed a bimodal distribution with peaks at 700 bp and 25 kb, while INVs (mean: 5.3 Mb, range: 54 bp to 118 Mb), exhibited a bimodal distribution centered around 1.5 kb and 2 Mb. These patterns are similar with prior study on somatic SVs in HCC(Zeng et al. 2025).

**Fig. 4.**
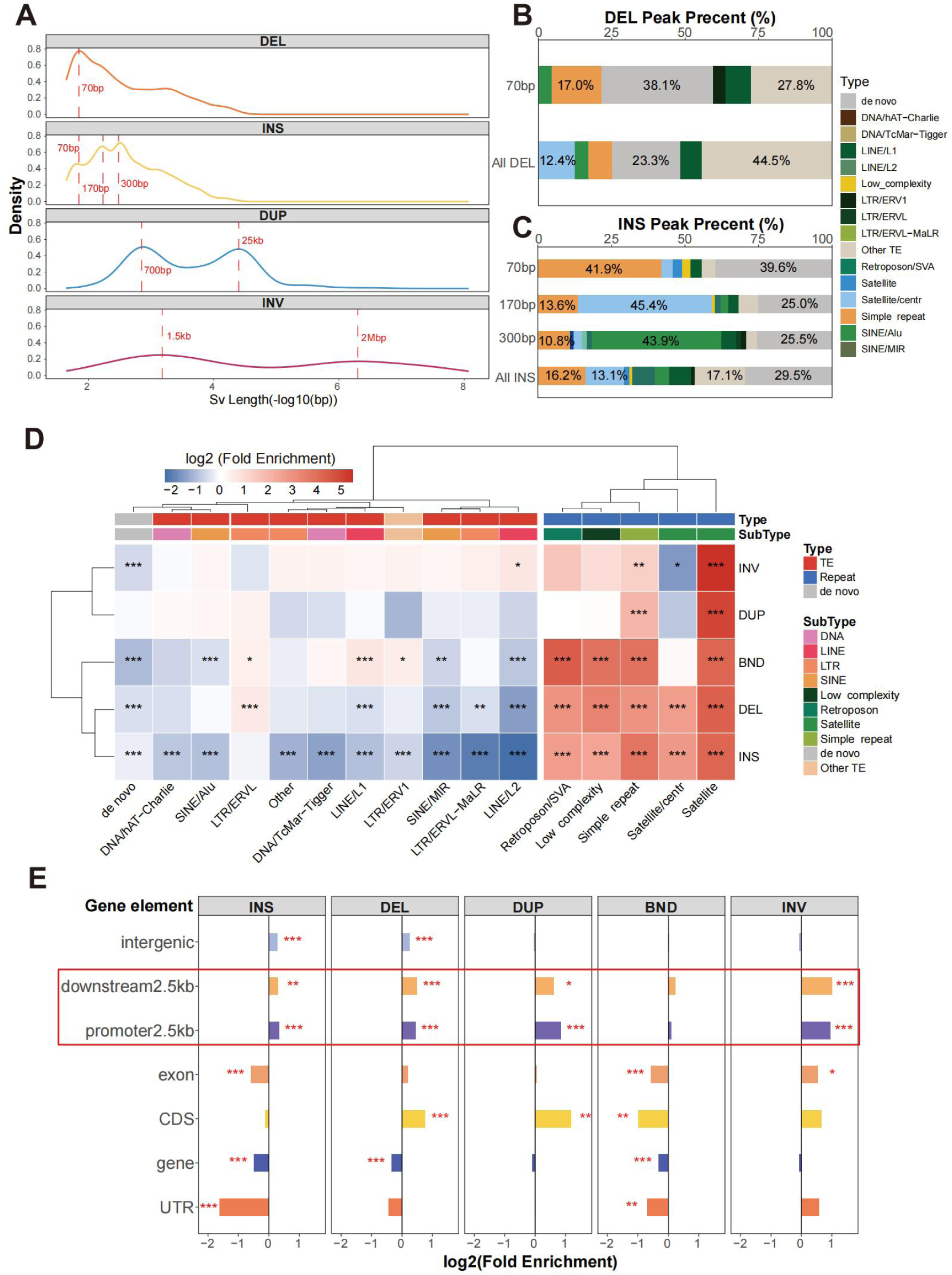
**Overview of SV length and site enrichment. (A).**Length distribution plot for different SV types. From top to bottom are DEL, INS, DUP, and INV, with red dashed lines marking the peaks for each SV type. **(B).**Stacked bar chart of sequence annotation proportions at DEL peaks.**(C).**Stacked bar chart of sequence annotation proportions at INS peaks. **(D).**Heatmap of somatic SV site enrichment in repeat regions. Each row represents a different SV type, and each column represents a different repeat region type, with color intensity indicating log2(Fold Enrichment). Repeat regions are categorized into TE (transposable elements region), Repeat (repeat region), and de novo (non repititive regions). **(E).**Bar chart of somatic SV site enrichment for gene elements. The X-axis represents log2(Fold Enrichment), the Y-axis represents gene element names, and facets represent different SV types. (*P < 0.05, **P < 0.01, ***P < 0.001)

To investigate the sequence determinants of somatic SVs, we analyzed the sequence feature of SVs at characteristic size peaks. Given the relatively uniform length distributions of DUPs and INVs (Fig. 4A and S4A), we focused on DELs and INSs (Fig. 4B and 4C). At the 70 bp peak, both somatic DELs and INSs were significant enriched in simple repeat sequences. Specifically, somatic DELs exhibited a 2.1-fold increase in simple repeats (17.0% vs 8.2% genome-wide somatic DELs, Pearson’s Chi-squared test, p=0.0956), while INSs showed a 2.6-fold enrichment (41.9% vs 16.2% genome-wide somatic INSs, Pearson’s Chi-squared test, p=0.00012). Notably, AAAG motifs predominated in somatic DEL (30%, Supplemental Fig. S4B), whereas somatic INSs exhibited a strong AT-richness (58.6% AT, Supplemental Fig. S4C). The 170-bp INS peak showed a notable association with centromeric regions. 45.4% of INSs harbored satellite/CEN (monomers: ∼170 bp(Aldrup-MacDonald et al. 2016), Pearson’s Chi-squared test, p=1.1e-6), with 55.6% localized within centromeric regions, compared to 12.3% of genome-wide INSs (Pearson’s Chi-squared test, p<0.001, Supplemental Fig. S4D). Interestingly, 170 bp INSs not located within centromeric regions tended to cluster near near centromeres (Wilcoxon test, p<0.001, Supplemental Fig. S4D), with 62.7% found within 1 Mb of centromeric regions (Wilcoxon test, p<0.001, Supplemental Fig. S4D), further emphasizing the variability of centromeric satellite sequences. At the 300-bp peak, SINE/Alu elements, primarily the AluY subclass, were predominant in INSs, comprising 43.9% of the total (Pearson’s Chi-squared test, p=3.0e-10) (Supplemental Fig. S4E). This observation may be attributed to the AluY subfamily, the youngest and most active member of the Alu family(Hormozdiari et al. 2011; Witherspoon et al. 2010), whose high activity has been identified in both cancer and healthy populations(Wu et al. 2021; Zeng et al. 2025; Ma and Pl 2002).

Somatic SVs breakpoints were non-uniformly distributed across the genome, with significant enriched in repetitive elements (TE enrichment score: 0.85; repeat enrichment score: 10.20; non-repetitive regions enrichment score: 0.82). Hyper-repetitive regions (including satellite, simple repeats, low complexity regions and SVA elements) showed substantial enrichment (mean enrichment score >6.56, Fisher’s test, p-value <0.05, Fig. 4D). Notably, somatic DELs and INSs were more frequently observed in centromeric satellite sequences (enrichment score >6.4, Fisher’s test, p-value <0.05), whereas larger SVs, such as INVs and DUPs, were underrepresented in these regions (enrichment score <0.54). This pattern may be due to satellite sequences’ susceptibility to replication fork slippage or mispairing, combined with the compact chromatin structure of centromeric regions that limits DNA accessibility and favors conserved repair mechanisms, reducing the occurrence of large-scale SVs(Kashi and King 2006). Sequence annotation revealed that over 60% of INSs inserted in satellite/centr, satellite, and simple repeat regions harbored inserted sequences identical to flanking repeats (Supplemental Fig. S4F). A representative case in simple repeats demonstrated a 12-fold amplification of the reference TTCT motif in tumor samples (Supplemental Fig. S4G), indicating that somatic INSs preferentially propagate existing repetitive architectures through replication- or repair-associated templated insertion. Futhermore, within TE regions, BNDs and DELs exhibited targeted disruption of specific TE families (enrichment score > 1.3, Fisher’s test, p-value <0.05, Fig. 4D). While DUPs showed some TE families preference, these associations lacked statistical significance. In contrast, INVs exhibited broad enrichment across TE families (8/10) (enrichment score >1.2), while INSs were systematically depleted in most TE families (9/10 families) (enrichment score <0.55, Fisher’s test, p-value <0.05).

To elucidate the regulatory consequences of somatic SVs, we performed an enrichment analysis of SV breakpoints across functional genomic regions. Somatic INS, DUP, DEL, and INV exhibited significant enrichment within proximal transcriptional regulatory domains (Fig. 4D), particularly in promoter-proximal (2.5kb) and termination regulatory regions (downstream 2.5 kb), suggesting that these SVs may alter gene expression through cis-regulatory interference(Cosenza et al. 2022). We identified a strong spatial association between gene-proximal SVs and cis-regulatory elements. Specifically, INS and BND breakpoints were broadly localized to cis-regulatory elements, with the strongest enrichment observed at CTCF-binding sites (enrichment score >3.19, Fisher’s test, p-value <0.05, Supplemental Fig. S4H), suggesting potential disruption of topologically associating domain boundary. In contrast, DUP and DEL breakpoints preferentially disrupted enhancers and promoters near transcription/termination sites, suggesting they prefered modify the regulatory relationships between elements and genes (enrichment score >1.40, Fisher’s test, p-value <0.05). INV breakpoints lacked significant regulatory element predilection.

### Identify Extensive Figh-frequency Somatic SVs Regions in LSCC

To detected recurrent somatic SV across genome, we performed a non-overlapping sliding window approach for sample carrier rate quantification. We identified 226 high-frequency SV regions (sample carrier rate > 26%) exhibiting non-random chromosomal distribution (Fig. 5), and significant co-localization with known tumor-related genes(Tate et al. 2019) (Chi-squared test, p < 0.05,Supplementary Table S4). A example involves recurrent ∼150 bp DEL within the intronic region of C4BPA (4 tumors shared), a gene encoding a protein involved in complement activation, which plays a crucial role in cancer progression and served as a biomarker for early detection of pancreatic cancer(Sogawa et al. 2021).

**Fig. 5.**
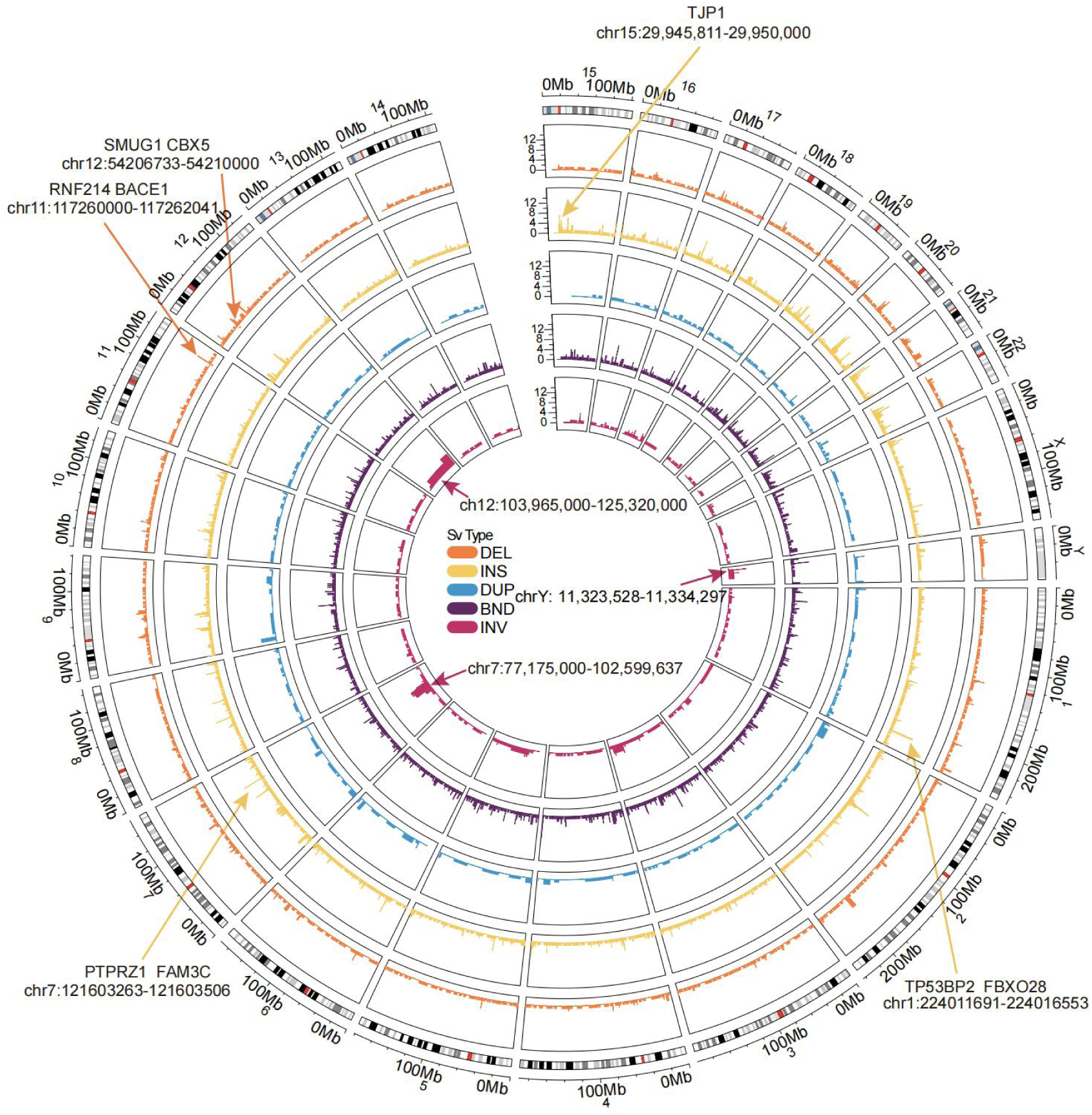
Circos plot of genome-wide sample enrichment. The tracks in the figure represent the enrichment of different SVs types in samples, from the innermost to the outermost being INV, BND, DUP, INS, and DEL. The plot highlights regions with the top 2 sample counts for INS and DEL that may affect genes, the top 3 regions for INV sample counts, and regions intersecting with those reported in previous studies.

Previous studies identified CNVs as key genomic rearrangements in HNSCC(The Cancer Genome Atlas Network 2015). Notably, 24.24% of CNV hotspots identified in the TCGA LSCC dataset co-localized with recurrent somatic SVs, indicating these regions exhibit high genomic instability or fragility. In CNV deletion regions, such as 2q36, 7q36.1, 16q24.2, and 18q1.22, more than five LSCC samples carried somatic DELs. Notably, in the 2q36.2 region, 11 samples harbored somatic DELs, three of which exceeded the CNV threshold (1 kb)(Redon et al. 2006) with lengths from 2,649 bp to 4,439 bp (Supplemental Fig. S5A). Similarly, in the 7q36.1 region, 11 samples carried DELs, two of which had lengths (LSCC12: 11,224 bp, LSCC14: 8,643 bp) exceeding 1 kb (Supplemental Fig. S5A). In contrast, in CNV amplification regions, 3q25.2 and 12p13.33, over four LSCC samples harbored somatic INSs/DUPs (Supplemental Fig. S5B). Specifically, in the 12p13.33 region, 14 samples carried INSs, seven of which had lengths (1,078 bp to 6,984 bp) exceeding 1 kb, and four samples had DUPs, three of which showed lengths (40,591 bp to 75,608 bp) far surpassing 1 kb. These results suggested that LRS offers a more detailed analysis of known LSCC CNV regions, revealing small insertions and deletions that may have been overlooked due to previous technical limitations.

Furthermore, we identified two high-frequency somatic INS regions, chr7:121,603,263–121,603,506 (shared by 9 LSCC samples, Supplemental Fig. S6A) and chr15:29,945,811–29,950,000 (shared by 7 LSCC samples, Supplemental Fig. S6B), which overlapped with recurrent repeat expansions (rREs) previously identified in the TCGA pan-cancer corhort(Erwin et al. 2023). The former INS aligned with a rRE specifically detected in lung squamous cell carcinoma, while the latter corresponds to a rRE characteristic of prostate cancers. The inserted sequences of these somatic INSs matched the motifs described in previous study, suggesting that these somatic INSs have the potential to serve as pan-cancer biomarker across multiple malignancies.

Moreover, we identified several high-frequency SVs that may influence nearby genes by disrupting regulatory elements. For instance, the region chr11:117,260,000–117,262,041 harbored somatic DELs (235-353 bp) in six tumors (Supplemental Fig. S6C). This region is located within an intron of *RNF214* and overlaped with an enhancer element predicted by ENCODE(Je et al. 2020). This enhancer is predicted to regulate *RNF214* expression in PC-3 prostate cancer cells(Je et al. 2020). *RNF214* is an E3 ubiquitin ligase. Its overexpression promotes tumor cell proliferation, migration, and invasion(Lin et al. 2024). Additionally, in chr12:54,206,733–54,210,000, somatic DELs (163-178 bp) in five samples overlapped with an enhancer predicted to regulate *SMUG1* expression in Panc1 pancreatic cancer cells(Je et al. 2020) (Supplemental Fig. S6D). *SMUG1* is involved in base excision repair and genomic stability maintenance(Iakovlev et al. 2019). High *SMUG1* expression correlates with poorer survival in the TCGA HNSCC cohort (Log-rank test, p = 0.003) (Supplemental Fig. S6E). We also identified eight somatic INVs shared by more than four samples (Supplemental Fig. S6F). The region chr7:77,175,000–102,599,637 showed the highest frequency (9 samples). Notably, 8 out of 9 samples harbored inversions exceeding 18 Mb, affecting multiple key genes, such as *CDK14*(Zhang et al. 2022) and *GNGT1*(Zhang et al. 2021). Large-scale inversions may disrupt gene architecture, leading to dysregulated expression of tumor-associated genes(Xu et al. 2023).

### Analysis and Validation of a Biomarker in 74.1% LSCC Patients

A somatic INS, observed in 66% (10/15) of LSCC tumors, was the most frequent recurrent SV in our study (Fig. 6A). It was located at chr1: 224,011,691-224,016,553, a simple repeat region annotated by RepeatMasker(Tarailo-Graovac and Chen 2009). The region initially consisted of a short tandem repeat with the (AATGG) repeat motif in normal samples, while the somatic alteration in 9/10 LSCC samples involved an expansion of this pattern (Fig. 6B).The LSCC11 sample differed, harboring Human Satellite II as its repeat motif (HSATII, a satellite DNA derived from the (GGTAA)n repeat)(Altemose et al. 2022). The somatic INS exhibited low supportting read ratios in six samples (10%-30%), while four exhibited stronger support (>50%) (Supplemental Fig. S7A). INS lengths varied among tumors (median: 581 bp, Fig. 6C and S7A). LSCC15 had a 3,610 bp INS, while other samples showed peak SRE lengths around 450 bp (mean 517 bp, SD 382 bp). Additionally, two patients harbored multiple INSs of different lengths. For example, the LSCC1 sample harbored 390 bp, 650 bp, and 1,030 bp INSs (Supplemental Fig. S7A and S7B), indicating more tumor heterogeneity.

**Fig. 6.**
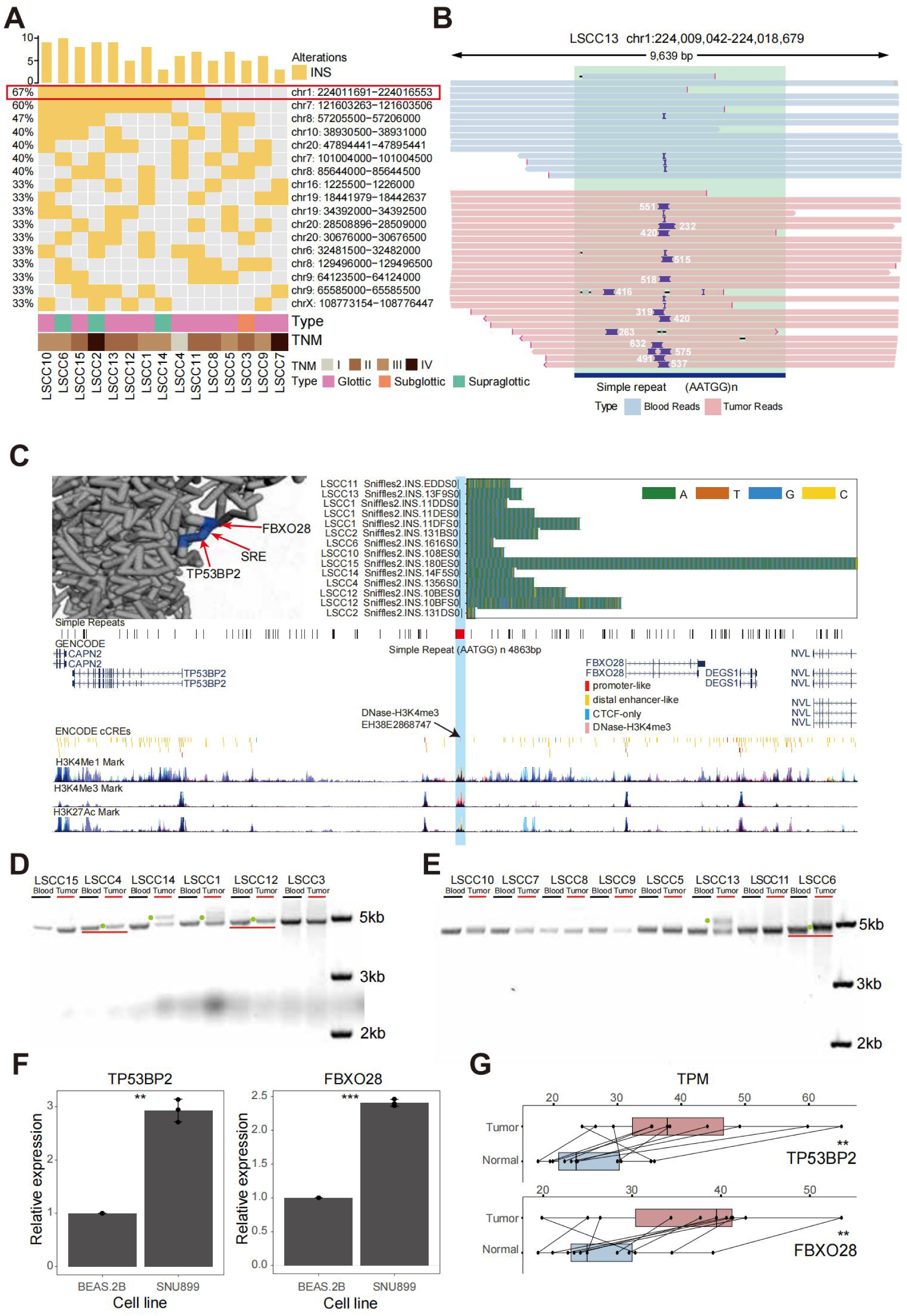
**Overview of somatic SREs shared by 66% of samples. (A).**Waterfall plot of INS sample enrichment regions. Rows represent samples, and columns represent regions. **(B).**IGV plot of somatic SRE (LSCC 13, chr1:224,009,042-224,018,679). Light blue represents reads from blood samples, pink represents reads from tumor samples, and dark blue indicates the Simple repeat region with the AATGG motif. **(C).**Browser plot of somatic SRE (chr1:224,009,042-224,018,679) illustrating the following epigenomic signals: a stacked bar chart of INS positions and inserted sequences, with different colors representing different bases; the plot concludes with the AATGG motif of the repeat, the ENCODE-annotated gene transcript, and Candidate Cis-Regulatory Elements (cCREs) predicted by ENCODE. Also included are H3K4Me1, H3K4Me3, and H3K27Ac marks predicted by ENCODE. In the upper left corner, the predicted 4D nucleosome structure around the LSCC-enriched SRE, TP53BP2, and FBXO28 is displayed. **(D).**Horizontal gel electrophoresis images 1 showing DNA fragments from blood and tumor tissues of LSCC samples sequenced using nanopore technology. Each sample’s blood and tumor are labeled separately, and green dots and red lines mark the longer bands in the tumor relative to the blood. **(E).**Horizontal gel electrophoresis images 2 showing DNA fragments from blood and tumor tissues of LSCC samples sequenced using nanopore technology. Each sample’s blood and tumor are labeled separately, and green dots and red lines mark the longer bands in the tumor relative to the blood. **(F).**Relative expression levels of TP53BP2 and FBXO28 determined by QCPCR in BEAS 2B and SNU899 cell lines. **(G).**Box plot of expression levels of TP53BP2 and FBXO28 genes in tumor and paired normal samples. Pink represents tumor samples, light blue represents paired normal samples, and P values were calculated by paired t-test. (*P < 0.05, **P < 0.01, ***P < 0.001)

To validated the presence of somatic INSs identified through LRS WGS, target PCR amplification of the INS region was performedon DNA from avialable 14 paired tumor and blood samples (including 9 tumors with INS identified by LRS WGS). Gel electrophoresis indicated that 6/9 tumors displayed longer fragments compared to blood samples (Fig. 6D and 6E). The remaining 3 samples presented challenges due to low INS supportting ratio (20% and 30% for LSCC10 and LSCC11) and a long insertion length in LSCC15 (3,610 bp)(Supplemental Fig. S7A), complicating the gel electrophoresis validation. Despite these challenges, our results confirm the authenticity and reliability of these high-frequency somatic INSs.

To assess the prevalence of the high-frequency somatic INS in LSCC tumors, we further collected 12 paired LSCC samples for PCR validation (Supplemental Fig. S7C-S7E, S8-S13). Gel electrophoresis and LRS data of the target PCR products revealed that 10/12 LSCC samples carried the somatic INS with an (AATGG) repeat motif, with insertion lengths ranging from 130 bp to 2,700 bp (Supplemental Fig. S8-S13). Furthermore, this INS was detected in the pericarcinomatous tissues of five LSCC tumors (Supplemental Fig. S10-S13). We also identified this INS in a lymph node metastasis sample from a patient with lymph node metastasis (Supplemental Fig. S14). Notably, the INS lengths in this patient’s tumor, lymph node metastasis, and pericarcinomatous tissuesshowed a reduction, measuring 400 bp, 200 bp, and 150 bp, respectively (Supplemental Fig. S9B-S9E). This observation suggested that this INS may confer a selective advantage during tumor progression and metastasis, potentially promoting tumor cell proliferation and migration, enabling its retention and expansion through clonal selection.

To further explore the functional role of the high-frequency somatic INS, we analysis the chromatin enviroment surrounding the INS. In the A673, A549, and SK-N-MC cell lines, the INS region overlapped with DNase-H3K4me3 signals, which are known to mark transcriptional start sites(Je et al. 2020)(Fig. 5C). While the target genes regulated by this DNase-H3K4me3 mark have not been extensively characterized, Hi-C data from the hypopharyngeal cancer FaDu cell line suggested that the region resides within the same topological domain (TAD) as the nearby coding genes *TP53BP2* and *FBXO28* (Supplemental Fig. S14A). Futhermore, data from the Four-Dimensional (4D) Nucleosome Consortium(Zhu et al. 2022) supported the presence of three-dimensional chromatin interactions between this DNase-H3K4me3 region and the genes *TP53BP2* and *FBXO28* (Fig. 5C). These findings raised the possibility that this INS may alter the regulation of these two genes. *TP53BP2* encodes the apoptosis-stimulating protein p53BP2, which interacts with p53 family members to modulate cell apoptosis and growth(Huo et al. 2023). *FBXO28*, which ecodes a ubiquitin ligase, has been reported to be aberrantly overexpressed in human epithelial cancer cell lines(Song et al. 2024).

We next examine the regulatory effect of the high-frequency somatic INS on gene expression. With gel electrophoresis, we identified longer fragments in two LSCC cell lines (SNU46 and SNU899) compared to a normal human bronchial epithelial cell line (BEAS-2B) (Supplemental Fig. S7C). Nanopore sequencing of the PCR products further confirmed that both LSCC cell lines harbored an expected INSs with AATGG repeat motif. Specifically, SNU46 and SNU899 carried homozygous INSs with lengths of 165 bp and 240 bp, respectively (Supplemental Fig. S14B-S14D). In contrast, no significant sequence length variation was observed in BEAS-2B cells compared to the reference genome, indicating the absence of the INS. We then selected the SNU899 cell line, which has a homozygous INS with longer insertion, and the BEAS-2B cell line for quantitative real-time PCR (QPCR) to assess whether the INS affects the expression levels of *TP53BP2* and *FBXO28*. We found that *TP53BP2* and *FBXO28* expression levels were significantly higher in the SNU899 cell line than in the BEAS-2B cell line (Fig. 6F, t-test, *TP53BP2*: p ≤ 0.01; *FBXO28*: p ≤ 0.001). In comparison, when examining gene expression in normal lung and larynx tissues, no significant upregulation of these two genes was observed in larynx tissue relative to lung tissue, indicating that the observed experssion differences was not tissue-specific (Supplemental Fig. S14E and S14F). This aligns with TCGA paired tumor-normal sample analysis, which showed significantly higher expression of both genes in LSCC tumor samples than in paired normal tissues (Fig. 6G, paired t-test, *TP53BP2*: p ≤ 0.01; *FBXO28*: p ≤ 0.01). Furthermore, QPCR analysis of tumor samples confirmed that those with the SRE exhibited significantly higher *TP53BP2* and *FBXO28* expression levels than those without the SRE (Supplemental Fig. S14G and S14H). These findings collectively supported the hypothesis that the high-frequency somatic INS upregulate the expression of *TP53BP2* and *FBXO28*, potentially playing a crucial role in the development and progression of LSCC.

## Discussion

Somatic structural variations (SVs) are critical hallmarks of tumorigenesis(Cosenza et al. 2022). However, technical limitations of SRS in resolving large-scale SVs, compounded by the tissue heterogeneity characteristic of HNSCC, have left the landscape of somatic SV in LSCC underexplored. This gap in knowledge has hindered the identification of LSCC-specific molecular biomarkers for effective cancer management. To address these challenges, we developed SomaGauss-SV, a LRS-optimized computational framework for somatic SV detection. Benchmarking against established tools revealed that SomaGauss-SV offered a superior balance of sensitivity and specificity, with an F1-score improvement of 11.3-33.9%. SomaGauss-SV excels in two key areas: (1) it reduces alignment errors in complex genomic regions, such as those surrounding rearrangements and repetitive sequences^59^, through a multi-step filtering approach, and (2) it accurately distinguishes true somatic INSs, DUPs and DELs in the presence of size-variable INSs/DUPs/DELs using a Gaussian mixture model. Despite these advances, significant technological challenges persist. Current LRS platforms still struggle with resolving complex breakpoint architectures, particularly those involving nested rearrangements and ultra-long SVs (>100 kb)(Keskus et al. 2025), limitations shared across existing detection tools, including SomaGauss-SV. However, recent innovations in high-throughput ultra-long-read sequencing technologies(Wang et al. 2021) and graph-based or assembly-based long-read alignment algorithms(Li and Durbin 2024) offer promise for overcoming these obstacles.

We systematically analyzed the LRS WGS data of 15 paired LSCC samples using SomaGauss-SV, revealing the distribution and sequence features of somatic SVs. We identified a dose-dependent positive correlation between smoking intensity and somatic DEL burden in LSCC. Previous studies indicated tobacco exposure is the primary driver of HNC, with tobacco users exhibiting higher frequencies of somatic SNVs and indels(Alexandrov et al. 2016; Torrens et al. 2025). Notably, tobacco-related mutational signatures show higher burdens and frequencies in laryngeal cancers compared to other HNC subsites(Alexandrov et al. 2016; Torrens et al. 2025). However, the link between tobacco and somatic SVs remains less clearly defined(Yoshida et al. 2020; Alexandrov et al. 2016; Torrens et al. 2025; Jethwa and Khariwala 2017). Our study fills this gap by demonstrating the role of chemical carcinogens in chromosomal instability. Specifically, we suggested that the accumulation of somatic DELs may result from the synergistic effects of multiple carcinogens in tobacco smoke. DNA adducts from polycyclic aromatic hydrocarbons and nitrosamines can block replication forks and cause DNA double-strand breaks(Yoshida et al. 2020). These breaks, when repaired poorly, may lead to chromosomal deletions. Also, free radicals and reactive oxygen species from tobacco can cause more DNA damage through oxidative stress, promoting deletions during faulty repair(Yoshida et al. 2020). Tobacco components might also disrupt cell division or cycle regulation(Aw et al. 2021), increasing chromosomal instability. Although high expression of *CYP1A1*, the key enzyme involved in the metabolism of benzopyrene and nitrosamines, has been proposed as a potential cause of the elevated tobacco-related mutation signatures in laryngeal tumors(Degawa et al. 1994; Hecht and Hatsukami 2022; Yamazaki et al. 1992), the specific tobacco carcinogens responsible for the observed DEL burden remain to be identified in vitro. Clinically, DEL burden could serve as a potential biomarker for risk stratification and therapeutic target screening in LSCC. Further investigation into high-frequency deletions associated with smoking may provide evidence for preventive interventions targeting high-dimensional genomic regions. For example, somatic DEL might disrupt CTCF binding in 3D chromatin architecture, influencing the expression of immune genes like *HLA-DRB5* and *HLA-DRA*. However, the limited sample size in this study restricts the generalizability of the observed association between somatic SV burden and smoking. Other potential confounders, such as alcohol consumption and dietary factors, may also influence the results. Future studies with larger corhorts and the incorporation of multivariable models or in vivo/vitro experiments are needed to confirm these findings.

In addition, 226 high-frequency SV loci were identified, exhibiting selective advantages during tumor clonal evolution. Cross-cancer comparison analysis revealed that high-frequency SVs at chr7:121,603,263-121,603,506 and chr15:29,945,811-29,950,000 overlap with hotspot somatic repeat expansions reported in pan-cancer studies(Erwin et al. 2023), suggesting their involvement in conserved carcinogenic pathways across various cancer types. Significant CNV regions in laryngeal cancer from the TCGA database also overlap with the high-frequency large DELs, DUPs and INSs, further validating the unique strength of LRS technology in resolving multi-scale genomic rearrangements. A key issue to explore further is how to define the boundary between CNVs identified through sequencing depth and DEL/INS/DUP detected by long reads.

Notably, analysis of both WGS and an independent validation cohort identified a somatic high-frequency SRE in 74.1% of LSCC patients. Out of 9 LSCC patients with adjacent cancerous tissue, somatic SREs were detected in 7 patients’ samples. In one patient, this SRE was detected across tumor, adjacent normal, and lymph-node-metastasis tissues, with lengths of 400 bp, 125bp, and 200bp, respectively. This suggested that the SRE evolves dynamically throughout tumor progression and metastasis, conferring varying selective advantages in different microenvironments. The longest SRE in the tumor likely formed early and stabilized to drive tumor growth. In contrast, the shorter SRE in the lymph-node-metastasis tissue may have undergone truncation but retained functionality, possibly reflecting adaptation to selective pressures associated with metastasis. The shortest SRE in the adjacent normal tissue could represent an early or ancestral form, implying that the SRE may pre-exist in peritumoral tissue and promote cell migration and colonization. However, the mechanisms linking SRE length variation to phenotypic diversity in tumor cells require further investigation.

The recurrent somatic SRE identified in this study holds significant potential as a biomarker for LSCC. While our study was limited by the insufficiency of paired multi-omics data, we hypothesized, through the integration of publicly available multi-omics data, that the SRE may regulate the expression of neighboring genes through a three-dimensional genome reconstruction mechanism. The STR region harboring the SRE exhibited abundant signals of H3K4me3, H3K4me1, and H3K27ac, suggesting the presence of sequences with promoter-like activity. We propose that the SRE-mediated elongation of this sequence could alter the spatial proximity between the distal promoter and the genes TP53BP2 and FBXO28, thereby enhancing their expression. Changes in this spatial interaction would require validation through 3C-PCR or Hi-C sequencing techniques. Given the tissue origin heterogeneity between the lung epithelial cell line BEAS-2B and the LSCC cell line SNU899, caution should be exercised in interpreting experimental results derived from these models. Previous studies have linked elevated FBXO28 expression with poor prognosis in ovarian cancer, where its upregulation activates the TGF-beta1/Smad2/3 signaling pathway, promoting cell viability, proliferation, migration, and invasion(Song et al. 2024). *TP53BP2*, a member of the ASPP (Apoptosis Stimulating Protein of p53) family, was reported to regulate apoptosis, proliferation, and autophagy, with its expression correlated strongly with cancer prognosis and patient survival(Huo et al. 2023). Notably, *TP53BP2* is highly expressed in human colorectal cancer (CRC) and promoted tumor formation or development, and conferring resistance to chemotherapeutic-induced apoptosis(Rieger et al. 2021). Therefore, further in-depth research on the molecular mechanisms of upregulated expression of *TP53BP2* and *FBXO28* in LSCC progression id crucial, and needed to be conducted using CRISPR-Cas9-edited cell models or conditional gene-edited animal models.

In conclusion, the bioinformatics pipeline developed in this study achieves an optimal balance between sensitivity and specificity, offering a robust tool for future research on tumor somatic SVs. This study represents the first comprehensive genome-wide survey of somatic SVs in LSCC using LRS, with the generated whole-genome sequencing data and identified SVs offering a valuable resource for clinical research in the field.

## Method

### Sample and Patient Information Collection

This study involved 27 patients with LSCC who underwent surgical resection at West China Hospital, Sichuan University. Tumor samples and matched blood samples were collected from these patients during surgery and venipuncture. Throughout the tissue collection and utilization process, this study strictly adhered to the Declaration of Helsinki and recieved formal approval from the Ethics Review Committee of West China Hospital, Sichuan University. All patients participating in this study have signed written informed consent forms. In addition, key clinical information, including clinical staging, smoking history, age, and gender, was throroughly documented for each patient.

### Cell Culture

Human cell lines, obtained from Procell (Wuhan Pricella Biotechnology Co., Ltd.) were utilized in this study. The cells were grown in DMEM medium (Gibco, C11995500BT) or RPMI 1640 medium (Gibco, C11875500BT), supplemented with 10% fetal bovine serum (FBS) and 100 U/mL Penicillin-Streptomycin (P/S). All cell lines were maintained at 37 °C and 5% CO2. Periodic mycoplasma testing was conducted using the Mycoplasma Detection Kit (Vazyme, D101). Cells were passaged at 70–90% confluency with 0.25% Trypsin-EDTA (Gibco, 25200-072).

### Real-Time Quantitative PCR

Total RNA was extracted using TRIzol reagent (Invitrogen, 15596026CN) and reverse-transcribed into cDNA with PrimeScript FAST RT reagent Kit (Takara, RR092A) following manufacturer’s protocols. Gene-specific primers (designed using NCBI and validated for specificity) were utilized with TB Green Premix Ex Taq II (Takara, RR820A) on a Bio-Rad system. All sequences were synthesized by Sangon (Chengdu, China) and are listed in the oligonucleotide table (Supplementary Table S5). Relative quantification was calculated using the method with GAPDH normalization.

### Validation of the SRE insertion in Tissues and Cell Lines using PCR and Sanger sequencing

Genomic DNA was isolated from tumor tissues, blood and LSCC cell lines using the QIAamp DNA Mini Kit (QIAGEN, 51306), followed by purity and concentration assessment via NanoDrop and agarose gel electrophoresis. The forward and reverse primers were designed according to the 500 bp upstream and downstream sequences of the insertion position and synthesized by Sangon Biotech (Supplementary Table S5). PCR amplification was performed in a Thermal Cycler using Phanta Max Super-Fidelity DNA Polymerase (Vazyme, P505).Each 50-μL reaction consisted of 2× Phanta Max Buffer 25 μL, 2 μL 10 μM forward primer, 2 μL 10 μM reverse primer, and 50-150 ng DNA. PCR was performed as follows: an initial step of 3 minutes at 95 °C, 30 cycles of 10 seconds at 95 °C, 15 seconds at 58 °C and 2 minutes 30 seconds at 72 °C. The PCR products were sent to Sangon Biotech for Sanger sequencing.

### Oxford Nanopore Library Preparation and Sequencing for HMW gDNA

High-molecular-weight (HMW) genomic DNA (gDNA) was extracted from tissue or blood samples using the MagAttract HMW DNA Kit (QIAGEN, 67563), and quantified using the Qubit® DNA Assay Kit and Qubit® 3.0 Fluorometer (Life Technologies, CA, USA). For library preparation, 9 samples (60%) were processed with the Ligation Sequencing Kit 1D (SQK-LSK109), and 6 samples (40%) with the Ligation sequencing DNA V14 kit (Oxford Nanopore Technologies, SQK-LSK114), following the manufacturer’s protocols (Supplementary Table S5). For the 60% of samples undergoing library preparation with SQK-LSK109, each sample was subjected to DNA end repair by combining 2 μg gDNA with 6.5 μL of the corresponding Buffer, 2 μL of NEBNext FFPE DNA Repair Mix (NEB, M6630L), and 3 μL of NEBNext Ultra II End Prep Enzyme Mix (NEB, E7546L), and the volume was adjusted to 60 μL with nuclease-free water. The mixture was gently mixed by pipetting, then incubated at 20 °C for 5 minutes followed by 65 °C for 5 minutes. The End-prepped DNA was purified with 1× volume of AMPure XP beads (Beckman Coulter, A63881) and washed twice with 80% ethanol. For adapter ligation, the purified DNA was mixed with 25 μL of Ligation Buffer, 10 μL of NEBNext Quick T4 DNA Ligase (NEB, E6056L) and 5 μL of Adapter Mix, then incubated at 25 °C for 20 minutes. Subsequently, 0.4× volume of AMPure XP beads was added, and the DNA pellet was resuspended in 25 μL of Elution Buffer. The remaining 40% of samples were prepared following the library preparation steps outlined in the SQK-LSK114 protocol. Libraries prepared with SQK-LSK109 were loaded onto R9.4.1 flow cells, while those prepared with SQK-LSK114 were loaded onto R10.4.1 flow cells. Sequencing was performed on the PromethION platform in accordance with the manufacturer’s instructions. Basecalling was conducted in batches using Guppy v3.2.8 with default parameters during sequencing.

### Oxford Nanopore Library Preparation and Sequencing for SRE PCR products

Fifteen SRE PCR products, amplified using Phanta Max Super-Fidelity DNA Polymerase (Vazyme, P505), were purified with AMPure XP Beads (Beckman Coulter, A63881) and quantified using the Qubit® DNA Assay Kit and Qubit® 3.0 Fluorometer (Life Technologies, CA, USA). For library construction of each sample, in a clean 0.2ml PCR tube, 400 ng of sample, corresponding buffer, 0.75 µL Ultra II End-prep Enzyme Mix (NEB, E7546L), and 0.5 µL NEBNext FFPE DNA Repair Mix (NEB, M6630L) were added, and the volume was brought to 15 µL with nuclease-free water. After gentle mixing and a brief spin, the reaction was incubated at 20 °C for 5 minutes and then at 65 °C for 5 minutes to complete end repair. Each end-prep reaction was purified with 1× volume of AMPure XP Beads (AXP) and washed twice with 80% ethanol. For barcode ligation, the purified sample was mixed with 2.5 µL Native Barcode (NB01-24), 10 µL Blunt/TA Ligase Master Mix (NEB, M0367L) in a 0.2ml PCR tube. After gentle mixing and spinning down, the reaction was incubated at room temperature for 20 minutes and then terminated with 2 µL EDTA. All barcoded samples were mixed and purified with 0.4× volume of AMPure XP Beads (AXP) and washed twice with 80% ethanol. For adapter ligation, the pooled barcoded sample was mixed with 5 µL Native Adapter (NA), 10 µL NEBNext Quick Ligation Reaction Buffer (5X), and 5 µL Quick T4 DNA Ligase (NEB, E6056L) in a 1.5 mL Eppendorf LoBind tube. After gentle mixing and a brief spin, the reaction was incubated at room temperature for 20 minutes. The reaction was purified with 20 µL resuspended AMPure XP Beads and washed with 125 µL Long Fragment Buffer (LFB). Finally, the library was eluted with 12 µL Elution Buffer (EB) to obtain the sequencing-ready library. We loaded 50 fmol of the library onto R10.4.1 flow cells (FLO-MIN114) and sequencing on the GridION X5 platform following the manufacturer’s instructions. Basecalling was performed in batches by ONT basecalling software version 7.1.4 with default parameters during sequencing.

### Library Preparation for Qitan Tech Nanopore Sequencing

Twenty-three SRE PCR products amplified using Phanta Max Super-Fidelity DNA Polymerase (Vazyme, P505) were purified with AMPure XP Beads (Beckman Coulter, A63881) and quantified using the Qubit® DNA Assay Kit and Qubit® 3.0 Fluorometer (Life Technologies, CA, USA). For end repair of the DNA, each 200 fmol sample was mixed with 2.5 μL of DNA repair buffer, 1 μL of DNA repair enzyme (DRM), 1 μL of end repair enzyme (EPM), and nuclease-free water to a final volume of 20 μL. The reaction mixture was incubated at 30 °C for 10 minutes, followed by 65 °C for 10 minutes. For barcode adapter ligation, the end-repaired DNA was combined with 2 μL of barcode, 7.5 μL of 4× ligation buffer, and 1.5 μL of DNA ligase, and the reaction was carried out at room temperature for 10 minutes. The reaction was then terminated with 2 μL of stop reaction buffer (TRB). All barcoded samples were pooled, purified with 0.4× volume of AMPure XP Beads, and washed twice with 80% ethanol. Subsequently, the sample was subjected to an additional purification step with 1× volume of AMPure XP Beads and eluted in 35 μL of nuclease-free water. For sequencing adapter ligation, in a 1.5 mL Eppendorf LoBind tube, 200 fmol of the purified product was mixed with 3.5 μL of sequencing adapter (SAC), 12.5 μL of 4× ligation buffer, 2 μL of DNA ligase (DLE), and nuclease-free water to a final volume of 50 μL. The reaction was performed at room temperature for 10 minutes, after which it was purified with 0.4× volume of AMPure XP Beads, washed with long fragment wash buffer (LWB), and finally eluted in 16 μL of elution buffer.

The final libraries were quantified via Qubit® 3.0 Fluorometer (Life Technologies, CA, USA) and loaded onto a QCell-384 V1.0 flow cell (Qitan Tech, Q-001-001-01). The sequencing was carried out using a QNome-3841 platform (Qitan Tech, China), strictly adhering to the manufacturer’s protocols. Base calling was performed using QPreasy v3.2.1 with default parameters in real time during sequencing.

### Long-read Whole-genome Sequencing Data Preprocessing

1. Fastq File Filtering

We performed quality filtering on the raw fastq files from both tumor and blood samples using NanoFilt software (version 2.8.0)(Lee et al. 2021), with parameters “-q 7 -l 1000”. The filtering criteria retained reads with a sequencing quality score greater than 7 and a sequence length greater than 1000 bp. Through this process, we obtained clean fastq files, which contained sequence data that met the quality standards.

1. Quality Control of Clean Fastq Files

After the initial filtering of the clean fastq files, we further conducted quality control using NanoPlot software (version 1.39.0)(De Coster and Rademakers 2023) with parameters “--maxlength 40000 --plots hex dot”. During this process, we filtered out reads shorter than 40,000 bp and performed a detailed statistical analysis on these reads, including calculating key sequencing information such as the N50 value and total base count.

1. Alignment to the Reference Genome

We used minimap2 (version 2.26)(Li 2018) to align the clean fastq files from both tumor and blood samples to the hg38 reference genome from NCBI, generating SAM files with parameters “-MD -ax map-ont -L”. Subsequently, we utilized Samtools (version 1.9)(Danecek et al. 2021) to convert the SAM files into BAM files and sorted the BAM files to obtain sorted BAM files (sort bam). Then, with the help of Samtools (version 1.9) and awk commands, we added “tumor” and “blood” tags to each read in the sort bam files for tumor and blood samples, respectively, ultimately generating sort tag bam files.

### Analysis and Visualization of Nanopore Sequencing Data for PCR Products

After nanopore sequencing of the PCR products, we utilized the NanoFilt software (version 2.8.0) with parameters “-q 7 -l 4800” to filter reads that could fully span the repeat regions. Subsequently, we aligned the filtered reads to the hg38 reference genome from NCBI using minimap2 (version 2.26), generating SAM files with parameters “-MD -ax map-ont -L”. Thereafter, we converted the SAM files to BAM files using Samtools (version 1.9). To avoid alignment errors, we excluded reads containing DEL longer than 50 bp through an in-house bioinformatics pipeline, considering that INS in such reads were likely due to alignment artifacts. Finally, we visualized the SRE-containing paired samples using the Integrative Genomics Viewer (IGV, version 2.16.0)(Robinson et al. 2011).

### Acquisition and Preprocessing of Nanopore Sequencing Data and Gold Standard Somatic Structural Variations Data from Cell Lines

According to the reference(Keskus et al. 2025), we downloaded the nanopore sequencing data for five paired cell lines: HCC1395/HCC1395BL, H1437/H1437BL, H2009/H2009BL, HCC1937/HCC1937BL, and HCC1954/HCC1954BL, as well as the corresponding gold standard sets of somatic structural variations for these cell lines. To ensure the accuracy of the analysis, we filtered the gold standard data, retaining only those somatic structural variations that could be detected by the nanopore sequencing platform, totaling 4058, including DEL 1493, INS 906, DUP 385, INV 48, and BND 1226. Subsequently, following the standard workflow for nanopore sequencing data preprocessing, we aligned and processed these data, ultimately obtaining BAM files with “blood” and “tumor” tags, respectively.

### SomaGauss-SV Workflow

1. Merging BAM Files and Detecting Structural Variations

Using Samtools (version 1.9), we merged the sorted tag BAM files of paired samples to generate a merged BAM file. Subsequently, we utilized Sniffles2 (version 2.0.6)(Smolka et al. 2024) to analyze the merged BAM file with parameters “-t 100 --minsupport 3 --mapq 50 --min-alignment-length 1000 --output-rnames --allow-overwrite --long-ins-length 100000 --cluster-binsize 500 --minsvlen 50”. We filtered SVs based on the following criteria: read alignment quality greater than 50, supporting read count exceeding 3, SV length greater than 50 bp, and minimum alignment length greater than 1000 bp. The filtered SV information were then recorded in the merge.vcf file.

1. Screening for Candidate Somatic Structural Variations

To accurately identify candidate somatic SVs, we implemented a multi-step screening strategy. First, an in-house script was used to preliminarily filter SVs reported by Sniffles2, retaining only those supported by reads tagged with “tumor”, which were initially considered as candidate somatic SVs.

Based on the candidate somatic SVs, we further refined the screening process for DELs as follows:

λ Region amplification and small BAM file generation: Using the start position of the candidate DEL as the center, we amplified a 500 bp region on both sides. Reads from these regions were extracted from the complete tumor and blood BAM files to generate corresponding small BAM files.

λ DEL detection and preliminary screening: Using the pysam tool, we scanned each read in these small BAM files to find deletion events greater than 50 bp in length and similar to the DEL length reported by Sniffles2 (difference less than 20 bp). If no DEL meeting these criteria was found in the blood BAM, the candidate DEL was automatically determined as a genuine somatic DEL.

λ Application of the mixed Gaussian distribution model: If DELs meeting the above conditions were found in the blood BAM, we input these DEL length data into the mixed Gaussian distribution model for analysis. We then distinguished DELs supported only by tumor reads (supporting reads greater than 3) and identified these as genuine somatic DELs.

λ Overfitting issue resolution: To address potential overfitting in the mixed Gaussian model, we introduced an additional Student’s t-test to compare the differences in DEL lengths between blood and tumor. Only when there was a significant difference in DEL lengths between blood and tumor, and the DEL was supported by with more than three tumor reads, was it ultimately identified as a genuine somatic DEL.

For candidate somatic INS/DUPs, we employed the following screening strategy:

λ Region amplification and re-detection: Using the start position of the candidate INS/DUP as the center, we amplified a 500 bp region on both sides to form a region list. INS variants in these regions were then re-detected using Straglr [10.1186/s13059-021-02447-3], with parameters “--genotype_in_size --min_support 1 --max_str_len 50 --nprocs 200 --min_ins_size 50 --max_num_clusters 9”.

λ Preliminary screening: If an INS/DUP greater than 50 bp in length was also detected in the same region in the blood sample, and this INS was grouped with the INS in the tumor, the candidate INS/DUP was excluded. Otherwise, it was retained. After this step, the remaining candidate INS/DUPs were identified as genuine somatic INS/DUPs.

Through this rigorous screening process, we effectively reduce false positive caused by alignment errors and Sniffles2 clustering algorithms, thereby improving the accuracy and reliability of somatic SV identification.

1. Normal Population Structural Variation Filtering

In the analysis of clinical sample pairs, we employed an additional screening step to further optimize somatic SV identification, which was not applied to cell line samples. Using Jasmine (version 1.1.5)(Kirsche et al. 2023), we wrote the absolute paths of the VCF files obtained from the previous step, as well as the absolute paths of the structural variation VCF files of the Chinese normal population, as reported in previous literature(Wu et al. 2021), into a txt file (named “sample-VCF.txt”) as input.cific steps are as follows:

λ Generating the out_vcf file: Jasmine was used to integrate the two VCF files, generating a new VCF file (named “out_vcf”) with parameters “threads=30 min_seq_id=0.9 k_jaccard=9 --dup_to_ins ignore_strand max_dist=500”.

λ Final screening and result determination: Further screening of the out_vcf file was performed using an in-house script. Specifically, SVs identical to those found in the Chinese normal population were removed, and the remaining variants were retained. These retained variations were ultimately identified as somatic structural variations in LSCC samples.

Through this series of steps, we can effectively distinguish somatic SVs in LSCC samples from common variations present in the normal population, thereby enhancing the accuracy and specificity of somatic SV identification.

1. Manual Screening for INV and BND

To enhance the precision of detecting INV and BND, we employed a visual analysis approach. Specifically, we utilized the Integrative Genomics Viewer (IGV) software(Thorvaldsdóttir et al. 2013) to generate images of INV and BND detected across multiple samples. These images provide an intuitive display of the variations’ positions in the genome, the distribution of reads involved, and the characteristics of the variations, facilitating detailed observation and comparison of these complex variations.

Through manual screening, we meticulously analyzed these IGV images. During the screening process, we relied on the characteristics of the variations (such as the number of supporting reads, alignment quality, and boundaries of the variations) and their commonalities across different samples to select high-confidence INV and BND from the images. This process ensures the accuracy and reliability of the detection results, avoiding false positives due to algorithmic errors or data noise.

By combining visualization with manual screening, we can more accurately identify and validate INV and BND variations, thus providing higher-quality data support for subsequent biological analyses.

### Detection of Somatic Structural Variations with nanomonsv

According to the nanomonsv user manual(Shiraishi et al. 2023), we processed the tumor bam and blood bam files obtained from the cell line’s nanopore sequencing data, and set the parameters “--single_bnd --use_racon --threads 200 --control_panel_prefix” for analysis. Ultimately, we obtained the VCF file of somatic SVs detected by Nanomonsv.

### Detection of Somatic SVs with Severus

According to the Severus usage instructions(Keskus et al. 2025), we processed the tumor bam and blood bam files obtained from the cell line’s nanopore sequencing data and applied the parameters “-t 30 --output-read-ids --vntr-bed $vntr-bed” for analysis. This resulted in the VCF file of somatic SVs detected by Severus. The vntr file was sourced from https://github.com/KolmogorovLab/Severus/blob/main/vntrs/human_GRCh38_no_alt_analysis_set.trf.bed.

### Detection of Somatic Structural Variations with SVision-pro

According to the SVision-prousage instructions(Wang et al. 2025), we processed the tumor bam and blood bam files obtained from the cell line’s nanopore sequencing data and applied specific parameters “--preset error-prone --min_supp 3 --max_sv_size 1000000 --sample_name sampleid−−processnum60−−detectmodesomatic−−modelpathsampleid --process_num 60 --detect_mode somatic --model_path sampleid−−processn um60−−detectm odesomatic−−modelp athmodel_liteunet_1 024_8_16_32_32_32.pth” for analysis. Ultimately, we obtained the VCF file of somatic SVs detected by Svision-pro. The model file was sourced from https://github.com/songbowang125/SVision-pro/tree/main/src/pre_process/model_lite unet_1024_8_16_32_32_32.pth.

### Evaluating Somatic Structural Variation Software

According to the method for SV assessment described in the reference literature(English et al. 2025, 2022), we chose to use Truvari as the assessment tool to compare the results of somatic SV detection with the gold standard. During the assessment process, we adopted the parameter settings “bench --pctseq 0” reported in the previous literature(Shiraishi et al. 2023; Wang et al. 2025; Lin et al. 2022).

### Correction and Annotation of Somatic INS Consensus Sequences

1. Correction of Consensus Sequences

To enhance the accuracy of somatic INS sequences, we initially constructed new somatic INS VCF files based on the VCF list of authentic somatic INS. Subsequently, we employed Iris (version 1.0.4)(Kirsche et al. 2023) to correct the consensus sequences provided by Sniffles2, with parameters set as “genome_buffer=1000 --keep_long_variants threads=20”. Iris optimizes the consensus sequences by integrating information from the genomic reference sequence, VCF files, and BAM files, thereby reducing sequence biases that may arise from sequencing or alignment errors.

1. (2) Annotation of Consensus Sequences

Building upon the corrected consensus sequences, we further annotated them using RepeatMasker (version 4.1.2)(Tarailo-Graovac and Chen 2009). RepeatMasker is a widely utilized tool capable of identifying and annotating repetitive sequences in the genome, such as transposons and satellite DNA. By annotating the corrected consensus sequences, we can better understand the origins and potential functional impacts of insertional variants. The parameters for RepeatMasker were set as “-species human -engine RMBlast -q -parallel 4”.

### Annotation of Somatic DEL Sequences

To further analyze the genomic context of somatic DELs, we employed bedtools (version 2.30.0)(Quinlan and Hall 2010) to perform intersection analysis between the filtered somatic DEL breakpoints and the BED file of the hg38 reference genome annotated by RepeatMasker. This process aims to determine whether the deletion regions overlap with repetitive sequence regions in the genome and to obtain corresponding annotation information. The specific steps are as follows:

1. Intersection Analysis

Utilizing the intersect function of Bedtools, we conducted an intersection analysis between the breakpoints of somatic DELs and the BED file of the hg38 reference genome annotated by RepeatMasker. This step identifies the overlapping parts between the deletion regions and repetitive sequence regions.

1. Annotation Information Merging

When a deletion region intersects with multiple annotated regions, to integrate these annotation informations, we used the “|” symbol to merge the annotation contents. For instance, if a deletion region overlaps with several different types of repetitive sequences (such as SINE elements and LINE1 elements), these annotation informations will be merged into “SINE|LINE1”.

### Statistical Analysis of Enrichment Site Regions

1. Data Acquisition: Retrieve BED files for gene elements and candidate cis-regulatory elements (CCRE) from the UCSC database(Cm et al. 2020).
2. Intersection Analysis: Employ Bedtools (version 2.30.0) to conduct an intersection analysis between the somatic SV loci of each sample and the aforementioned BED files. Summarize and tally the number of somatic SV loci within each element and the length of these elements.
3. Calculation of Enrichment Fold and Significance: Using the total count of somatic SV s and the genome’s total length, compute the enrichment fold.
4. Chi-Square Test: With the aid of the fisher_exact package in Python, construct a contingency table based on the total number of somatic SVs, variations within elements, variations outside elements, and the genomic background. Perform a chi-square test to determine if there is a significant enrichment of sites in that region.

### Sample Enrichment Analysis

Based on previous literature reports(Sahlin et al. 2023; Ahsan et al. 2023) and the visualization of alignment data (Supplemental Fig. S1), we observed that alignment in repetitive regions, such as simple repeat sequences, may lead to ambiguity. Despite using relevant software to correct for these areas, accurately calculating somatic SV loci within repetitive regions remains challenging. To address this issue, we created non-uniform window BED files specifically for sample enrichment statistics.

1. Creation of Non-Uniform Window BED Files The specific process is as follows:

1. Extract simple repeat regions: Extract regions annotated as “Simple repeat” from the RepeatMasker annotation file of the reference genome.
2. Segment the entire genome: Utilize these “Simple repeat” regions to segment the entire genome.
3. Divide non-repetitive regions: Segment the genome, excluding “Simple repeat” regions, into windows of 500 bp/5 kb.
4. Merge BED files: Combine the BED files of the “Simple repeat” regions with those of the evenly divided windows to obtain the final non-uniform window BED file.
2. Enrichment Statistics of Somatic Structural Variations

1. Intersection analysis: Use Bedtools (version 2.30.0) to perform intersection analysis between the somatic structural variations of each sample and the non-uniform window BED files created above.
2. Summary statistics: Aggregate and count the samples and the number of samples present in each region to assess the enrichment of somatic structural variations across different regions.

By employing this method, we can more accurately determine the distribution of somatic SVs in both repetitive and non-repetitive regions, providing more reliable data support for subsequent biological analyses.

## Data availability

All LR WGS data are available at the GSA Human database https://ngdc.cncb.ac.cn/gsa-human/s/Y4Sv2ZB2 (accession number HRA008222). The LRS WGS data and benchmark SV sets of five cell line pairs were downloaded from NCBI SRA BioProject PRJNA1086849.

## Code availability

SomaGauss-SV is available at https://github.com/XuYan000131/SomaGauss-SV.

## Acknowledgements

We thank Ranlei Wei for managing the computational resources required for this study. This work was supported by grants from the National Natural Science Foundation of China (82173383 to DX, 32200508 to LX, 82371883 and 12441517 to JFL), the National Key Science and Technology Special Project for Deep Earth Research (2024ZD1000600 and 2024ZD1000606 to JFL), Foundation of Sichuan Provincial Science and Technology Program (2025ZNSFSC0728), the 1·3·5 project for disciplines of excellence, West China Hospital, Sichuan University (ZYYC23024 to D. Xie).

## Authors’ contributions

LX and XYL conceived and developed the methodology, performed bioinformatic analyses. JFL and YXQ collected the clinical samples and clinical data. BYY, XL, XY, XC and HHZ collected the clinical samples. YH and YL designed and performed experiments. LX, XYL, YXQ, YH wrote the manuscript. DX and JFL responsible for supervision of research, data interpretation, and manuscript preparation. All authors read and approved the final manuscript.

## Declarations

Ethics approval and consent to participate

This study utilized publicly available data from The Cancer Genome Atlas (TCGA) and the Genotype-Tissue Expression (GTEx) project. All samples in these databases were collected with patient consent and appropriate ethical approval from the relevant institutional review boards. Our study did not involve additional human participants, human data or tissue.

## Competing interests

Dan Xie is the co-founder of Qitan Technology. Other authors declare no competing interests.

## Notes

### Competing Interest Statement

The authors have declared no competing interest.

